# Temporal Regulation of Organelle Biogenesis

**DOI:** 10.1101/2022.12.30.522312

**Authors:** Smruti Dixit, N Aniruddha, Sandeep Choubey

## Abstract

Organelle abundance is tightly regulated in eukaryotic cells in response to external stimuli. The underlying mechanisms responsible for this regulation remains less understood. Time-lapse imaging of fluorescently labelled organelles allow for counting individual organelle copies at a single-cell level. These experiments contain information about the time evolution of distribution of organelle number across a population of cells. To tap onto such data, we build upon a recently proposed kinetic model of organelle biogenesis that explicitly incorporates de novo synthesis, fission, fusion and degradation of organelles. Different limits of this general model correspond to distinct mechanisms of organelle biogenesis. We compute the first two moments of organelle number distribution for these different mechanisms. Interestingly, different mechanisms of biogenesis lead to qualitatively distinct temporal behavior of cell-to-cell variability (noise), thereby allowing us to discern between these mechanisms. Notably, noise in temporal organelle abundance exhibits strikingly more complex behaviour as compared to the steady state. Our modeling framework paves the way for extracting quantitative information about the dynamics of organelle biogenesis from time-lapse experiments that can measure organelle abundance at single-molecule resolution.

## 1 Introduction

Eukaryotic cells consist of various membrane-bound organelles that carry out distinct functions. For instance, mitochondria are responsible for producing ATP, Lysosomes function as the cell’s recycling center, etc [1]. Cells regulate organelle abundance in response to environmental and intracellular cues [2]. Mitochondria number is regulated as a function of the metabolic needs of the cell [3]. Algal cells tightly regulate the biogenesis of chloroplasts as a function of the existing number of chloroplasts. How a cell regulates its organelle abundance remains elusive.

Live and fixed cell imaging techniques have been employed to uncover mechanisms of organelle biogenesis. These studies have identified various processes that regulate organelle number, namely, de novo synthesis, fission, fusion. Importantly, these processes are inherently stochastic, which leads to cell-to-cell variability in their abundance. One approach to unraveling the mechanisms of organelle biogenesis has been to count organelle number across a population of cells. The measured steady state distribution of organelles is then used to infer the mechanism of organelle biogenesis. In particular, Mukherji et al. [4] constructed a simple model of biogenesis involving the four aforementioned processes. By employing a combination of theory and experiments, they showed that different mechanisms of organelle biogenesis lead to distinct footprint at the level of their cell-to-cell variability. These distinct signatures allowed for discriminating between different mechanisms of organelle biogenesis. For instance, Golgi body biogenesis involves de novo synthesis and decay, which is characterized by a Poisson distribution. In contrast, vacuole abundance is controlled via fission and fusion, which led to a truncated Poisson distribution. Subsequent studies furthered this line of investigation [5, 6].

However, these studies operated under the assumption that organelle abundance distribution has attained steady state, whereby the distribution has ceased to change in time. While the assumption of steady-state has been useful in explaining experimental data, it is limited in its applicability. For instance, during cell division organelles are partitioned between daughter cells. Such partitioning alters the average organelle abundance as well as its cell-to-cell variability in the daughter cells. Hence, these cells need to attain steady state level to be functionally viable. Moreover, real time imaging of fluorescently labeled individual organelles at a single-cell level across a population of cells provide information about the temporal evolution of organelle distribution. Any theoretical understanding of the temporal evolution of organelle abundance that can connect mechanisms of organelle biogenesis to live imaging data is lacking. In this manuscript, we attempt such a task.

To this end, we employ a simple model of organelle biogenesis, as proposed earlier [4, 7]. We study the transient properties of the model. In particular, we consider a population of cells with a distribution of organelle abundance at a given instant and explore how this distribution evolves in time, as shown in Fig. 1.B. For various subsets of the model, the transient mean and variance are computed. We show that the transient properties of these two moments exhibit distinct behavior which allows us to discriminate between them. In particular, we show that, while analyzing fixed cell imaging data, conclusions drawn from the assumption of steady state can lead to misleading results. Overall, our study provides the necessary toolkit to analyze live-cell time-lapse imaging data.

**Figure 1:**
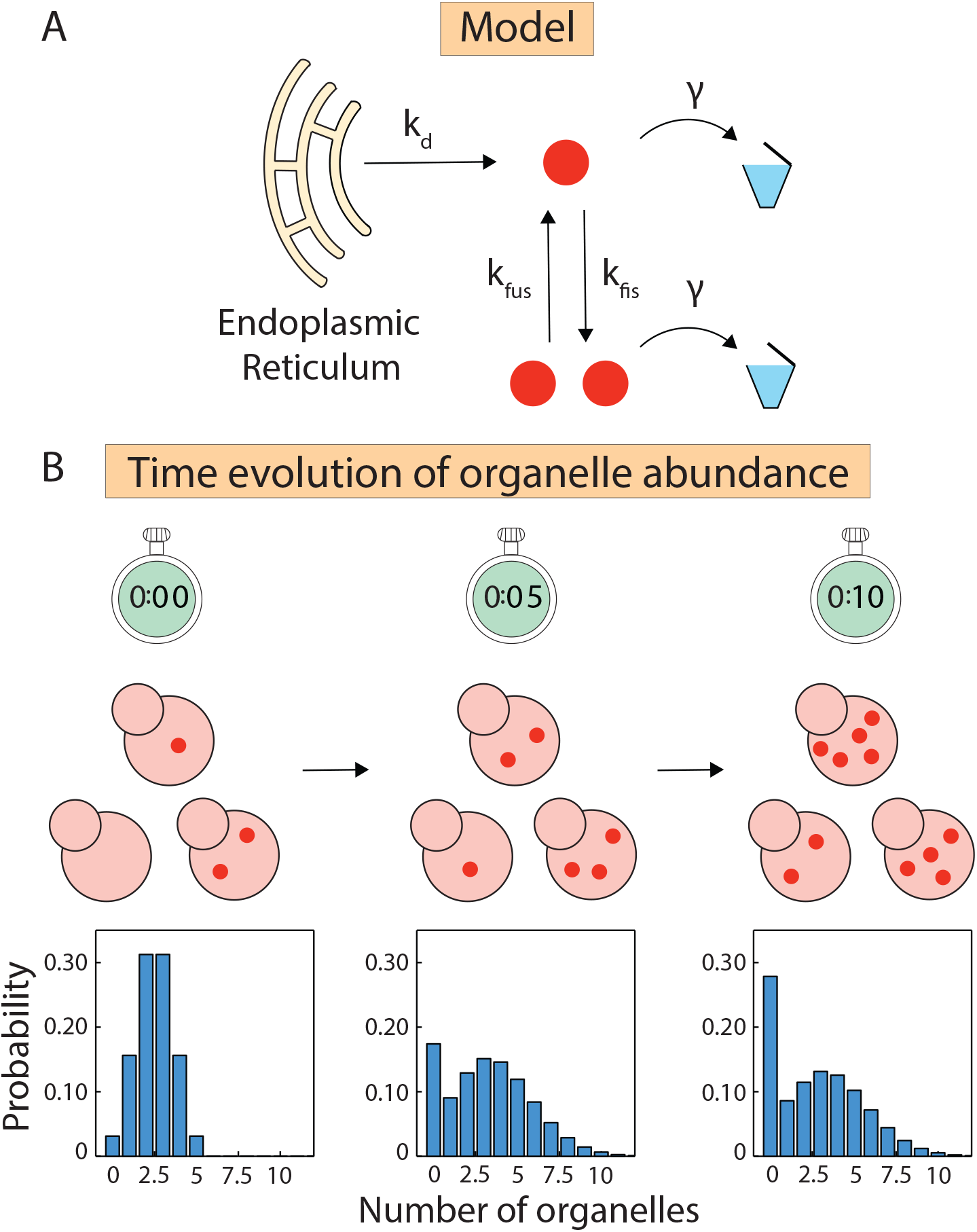
Model of organelle biogenesis. (A) We consider four processes which together govern organelle abundance in cells; these processes are de novo synthesis (*k_d_*), decay (*γ*), fission (*kfi*_s_) and fusion (*k*fus). (B) Live-cell imaging allows for monitoring organelle dynamics within a cell. Here we show a graphical representation of time-lapse snapshots of fluorescently labelled organelle copies in yeast. The bottom panel shows the time evolution of the histogram of organelle abundance across an isogenic cell population.

## 2 Model

To study the time evolution of organelle abundance, we employ a general model of organelle biogenesis, as proposed by Mukherji et al. [4] and used in subsequent studies. This model consists of four different processes: de novo synthesis, fission, fusion, and decay, as shown in Figure.1. de novo synthesis of an organelle happens from the Endoplasmic Reticulum with a zeroth order rate constant *k_d_*. An organelle copy can undergo fission with a first order rate constant *k_fis_*, thereby producing two copies. Two organelle copies fuse at a rate *k*_fus_. Finally, an organelle copy can decay with a first order rate constant *γ*. While de novo synthesis and fission increase the number of organelles, fusion and decay decrease their number. Through an interplay between these opposing processes, the organelle abundance evolves in time, and eventually attains steady state. Here, we are interested in the transient behavior of organelle abundance.

To explore the transient properties of the model, we employ a master equation-based approach, and monitor the time evolution of the organelle number. The probability *P_n_*(*t*) of having *n* organelles in a cell at a time *t* is given by the master equation

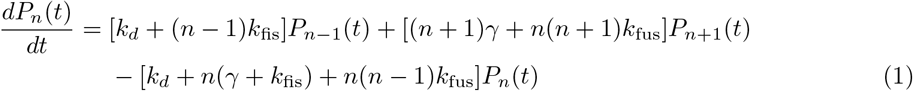

This equation is a collection of all possible ways that lead to either an increase or decrease in organelle copy number. Using this equation, we compute the mean and variance of the transient distribution of organelle abundance. These moments allow us to discern between different models of organelle biogenesis.

## 3 Materials and Methods

### 3.1 Numerical method

We solve Eq. 1, to obtain transient solutions for organelle numbers. Due to the presence of the non-linear terms in *n*, corresponding to the fusion process, the solution to the master equation is not analytically tractable. Therefore, we use the method given by Smith and Shahrezaei [8] to solve a general, one-dimensional, one-step master equation. The original master equation is a set of infinite coupled differential equations, which we truncate at some large integer *τ*. The set of equations can then be written in matrix notation as

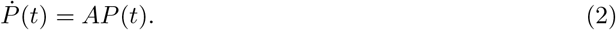

Here *A* is the transition matrix, given as

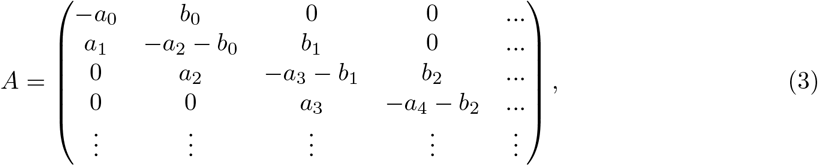

where *a_n_* = *k_d_* + (*n* – 1)*k*_fis_ and *b_n_* = (*n* + 1)*γ* + *n*(*n* + 1)*k*_fus_. Solving Eq. 2 requires evaluating the eigenvalues and the eigenvectors of the transition matrix *A*, which can become computationally intensive. Instead, using the fact that *A* is a tridiagonal matrix, and that the sum of each column of *A* is zero, the problem can be reduced to that of only evaluating its eigenvalues. Following the method given by Smith and Shahrezaei [8], we can numerically compute the transient distribution of organelle abundance. For a complete description, see the Supplementary information (SI). The transient probability distribution, evaluated in this manner, is given by

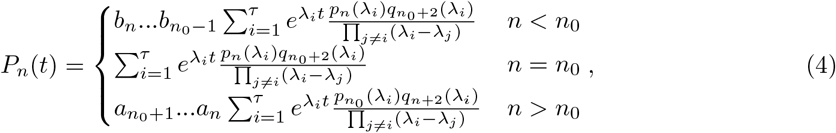

where *n*_0_ is the initial number of organelles, *λ_i_* is the *i*^th^ eigenvalue, and *p* and *q* are orthogonal polynomials defined recursively as

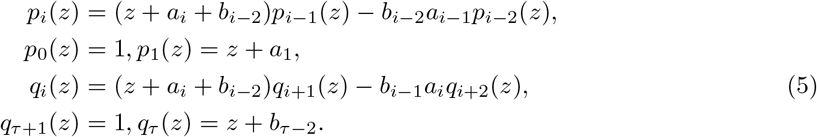

### 3.2 Analytical Method

The models without fusion do not contain the non-linear terms in *n*, and hence can be solved analytically. We use the generating function technique [9] to solve the set of coupled linear differential equations. Below, we describe the procedure briefly; see the SI for a more comprehensive description. The generating function is defined as

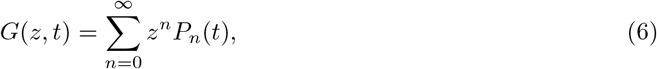

where 0 < *z* ≤ 1. Taking its derivatives with respect to *t* and *z*, and using them in Eq. 1, with *k*_fus_ and *k*_fis_ set to 0, gives us the following partial differential equation

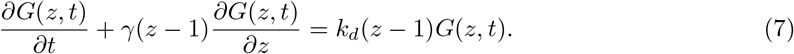

We can obtain the generating function by solving Eq. S14 using the method of characteristics [10]. The generating function is

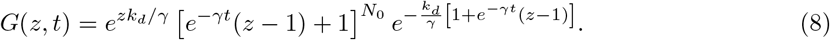

The moments of *P_n_*(*t*) can be obtained by taking the derivatives of the generating function, as follows

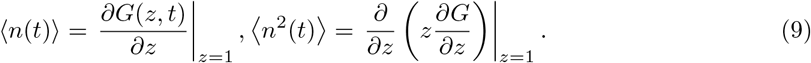

We use these numerical and analytical results to discriminate between the different models we consider.

### 3.3 Initial Conditions

The time evolution of the mean and Fano factor of organelle abundance is highly dependent on the initial conditions. We assume that initially the number of organelles across a cell population is binomially distributed, predicted by stochastic partitioning of organelles between daughter cells without any further regulatory control [11, 12]. However, experimental studies have shown that partitioning may not be random for some organelles, such as in the case of chromosomes or centrosomes where partitioning is strictly symmetric, with the daughter cells each inheriting one copy, whereas in endosomes and mitochondria, partitioning may be asymmetric.[13–15]. While we use binomial distribution of organelles as the initial distribution, we can easily compute the transient moments of organelle distribution for any of these aforementioned distribution of organelles.

## 4 Results

### 4.1 Mechanisms of organelle biogenesis govern the transient cell-to-cell variability in organelle abundance

To unravel the effect of different mechanisms of organelle biogenesis on the transient properties in organelle abundance, we consider different limits of the general model. Following up on previous studies [7], we consider five such limiting scenarios that have a corresponding finite steady-state, namely: (i) de novo synthesis-decay, (ii) fission-fusion, (iii) de novo synthesis-fission-decay, (iv) de novo synthesis-fusion-decay, and (v) de novo synthesis-fission-fusion-decay. For a detailed discussion on the limiting models, see the SI.

The master equation describing the time evolution of organelle numbers is one-dimensional. For such a system, the steady-state solutions for the mean, variance, and its distribution can be computed using different methods [4, 7, 16]. However, the presence of the fusion process renders the transient solutions inaccessible, owing to the corresponding nonlinear terms in the master equation. For models without fusion, i.e. (i) de novo synthesis-decay, and (ii) de novo synthesis-fission-decay, we compute the exact mean and variance of organelle distribution as a function of time using the generating function technique. For the rest of the models, we employ a recently developed numerical method by Smith and Shahrezaei [8] to compute the transient organelle distributions for various models. To find the details of the calculations, see the Methods and SI. In order to compare the cell-to-cell variability in organelle abundance for various models, we employ a statistical measure of dispersion, called the Fano factor. Fano factor is defined as the ratio of variance to mean.

For transient solutions, the initial condition needs to be specified. Here we consider two different scenarios. We assume the number of organelle copies across a population is binomially distributed. This initial condition is motivated by our question of how organelle abundance attains steady state after cell division. During cell division, the number of organelles is partitioned between the two daughter cells. For a random partitioning of organelles, the number of organelle copies in daughter cells will follow binomial statistics. This can be thought of as a coin toss problem, whereby each organelle copy is assigned to one of the daughter cells with a given probability.

In particular, we consider different limits of the general model and make specific predictions about transient organelle abundance as we tune the relevant model parameters.

#### 4.1.1 De novo synthesis-decay

First, we consider the de novo synthesis-decay model. Owing to its simplicity, this model provides a good starting point. This model has been shown to effectively capture the biogenesis of Golgi body [4, 17–20]. Since the steady-state organelle abundance is Poisson distributed, the Fano factor is one. We seek to investigate how the mean and Fano factor of organelle abundance approaches the steady state for this model. Using the generating function technique [9], we compute the mean 〈*n*(*t*)〉 and Fano factor *F*(*t*) as a function of time (see Materials and Methods and SI for details).

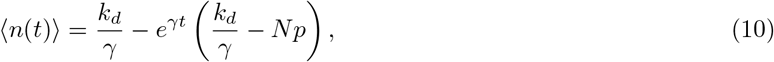

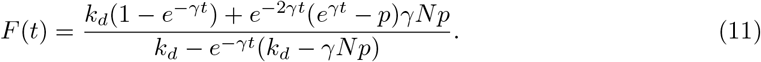

Using these analytical expressions, we find that the mean and Fano factor monotonically approach unity for parameter values and initial distributions (see SI Fig. S1), as shown in Fig. 4 A, B. These results suggest that the de novo synthesis-decay model can act as a reference to interpret the outcomes of more complex models. The rate of convergence of this model is dependant on *γ*.

**Figure 2:**
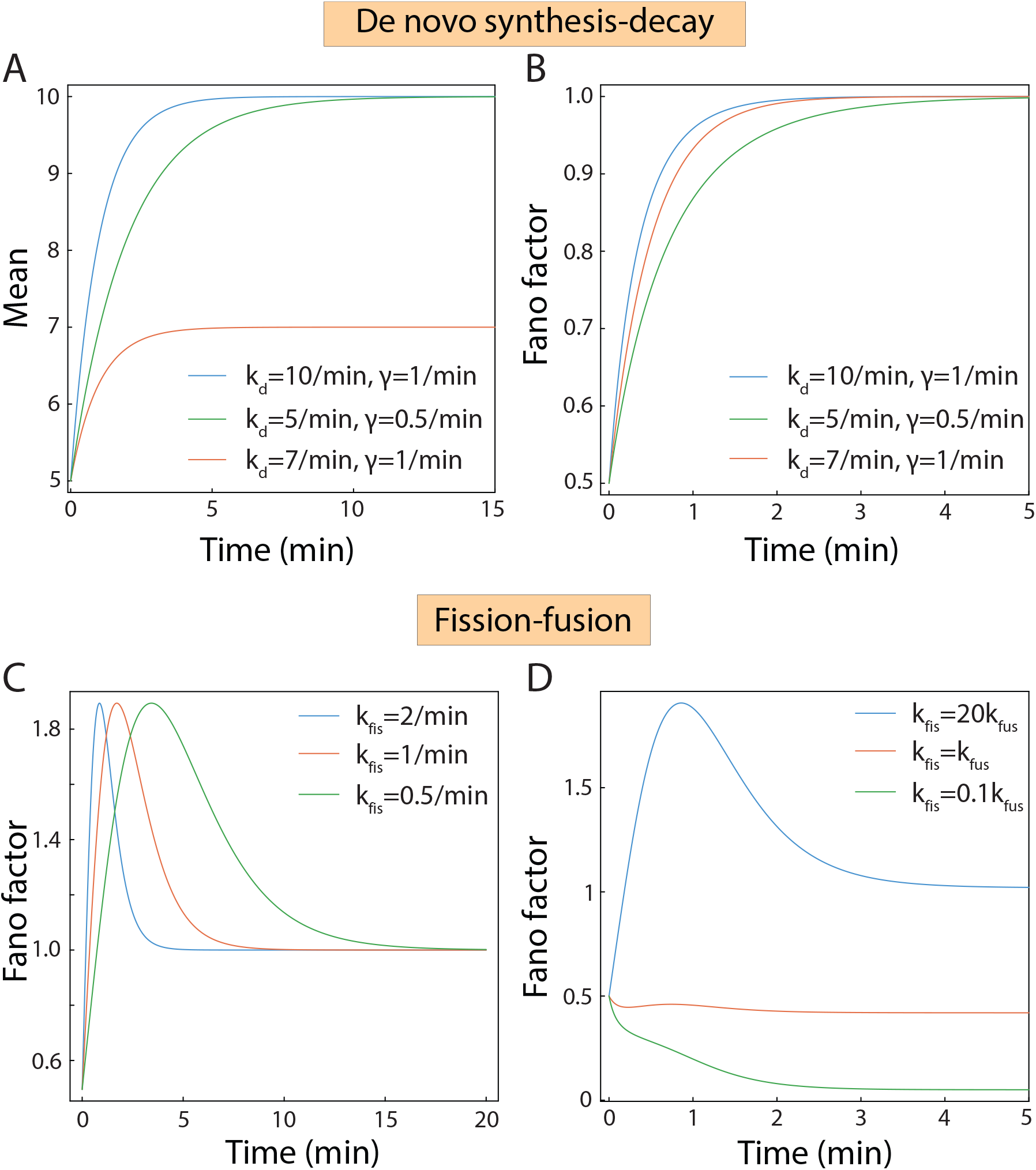
Mean and Fano factor profiles for various models. (A) De novo synthesis-decay: The mean number of organelles for various values of de novo synthesis and decay rate are shown. (B) De novo synthesis-decay: Here, we plot the time evolution of Fano factor for various values of *k_d_*; with *γ* = 1 min^-1^. (C) Fission-fusion: Dependence of the Fano factor keeping the ratio of parameters constant, *k*_fis_/*k*_fus_ = 20, and (D) Fano factor profiles for various values of fission rate *k*_fis_ and the fusion rate *k*_fus_

**Figure 3:**
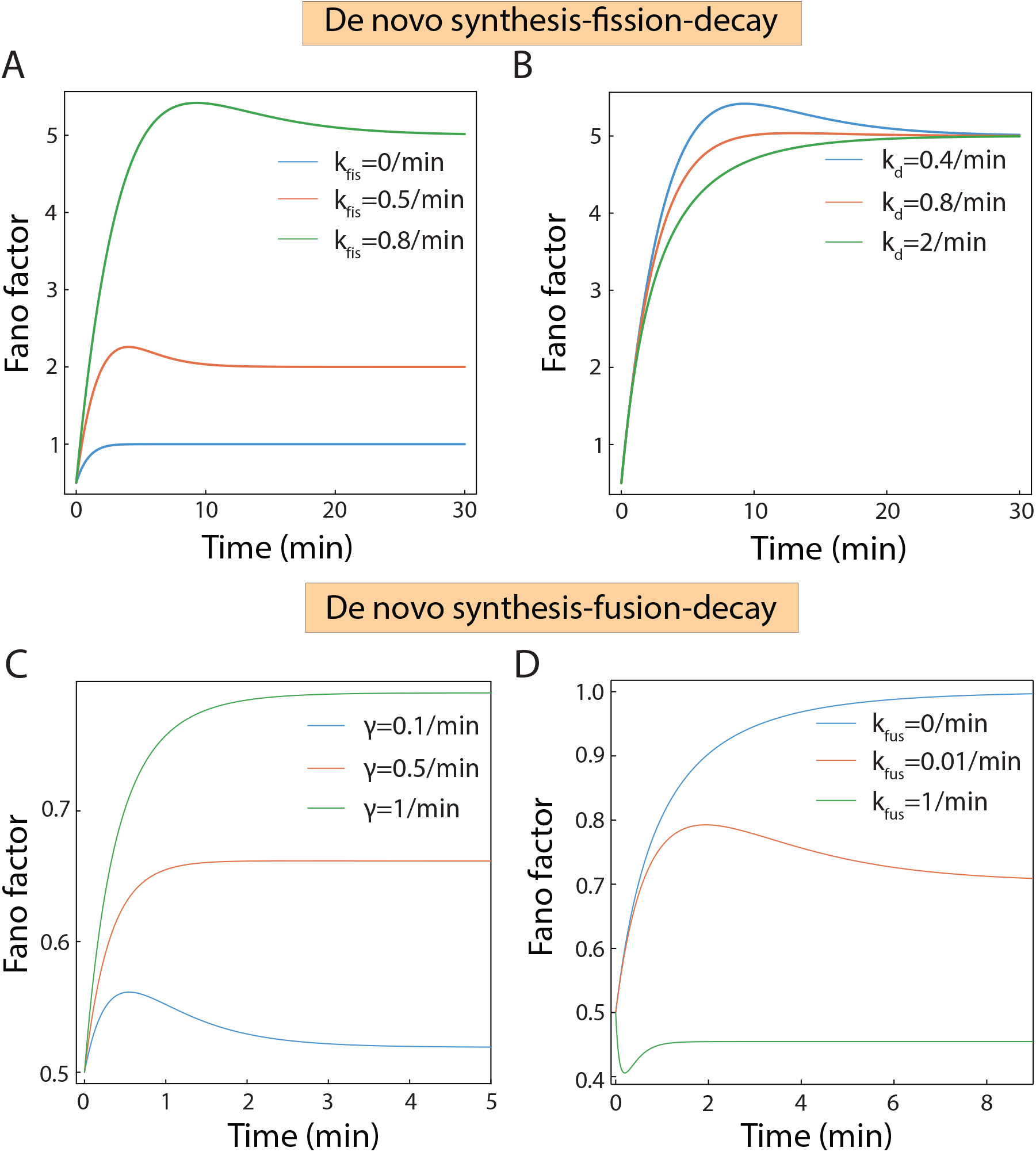
Time evolution of noise profiles for various models, with the initial number of organelles following a binomial distribution. (A) De novo synthesis-fission-decay: In this plot, we vary the fission rate *k*_fis_, while keeping the other rates constant; *k_d_* = 0.4 min^-1^, *γ* = 1.0 min^-1^. (B) De novo synthesis-fission-decay: Fano factor profiles for different values of the de novo synthesis rate *k_d_*, keeping *γ* =1 min^-1^ and *k*_fis_ = 0.8 min^-1^. (C) De novo synthesis-fusion-decay: Here, we plot the Fano factor as a function of time for various values of the decay rate *γ*, with *k_d_* = 5 min^-1^, *k*_fus_ = 0.1 min^-1^, and in (D) we vary the fusion rate *k*_fus_, with *k_d_* = 5 min^-1^ and *γ* = 0.2 min^-1^.

**Figure 4:**
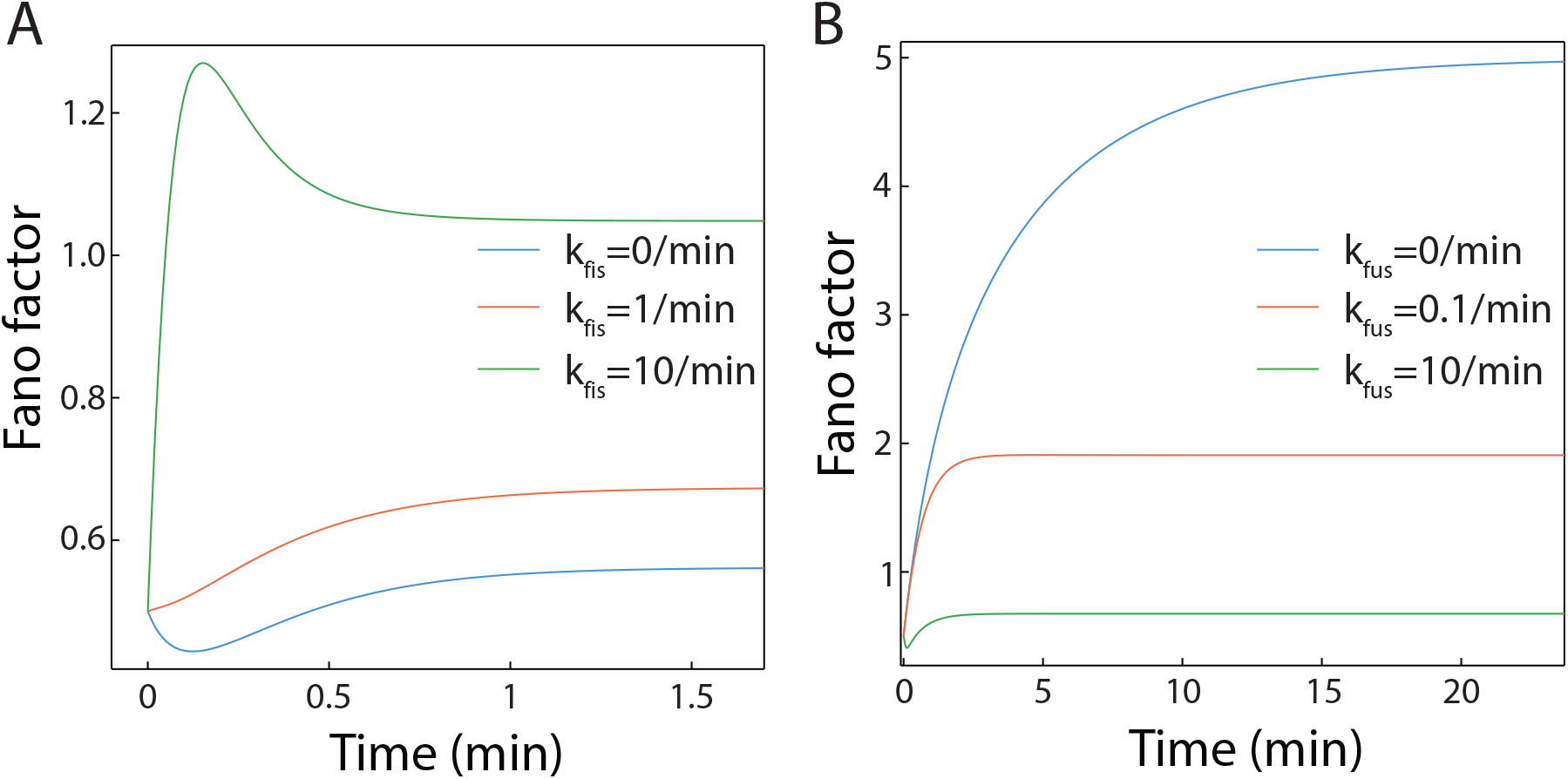
De novo synthesis-fission-fusion-decay. (A) In this plot, we vary the fission rate *k*_fis_ and look at the time evolution of the Fano factor, with *k_d_* = 5.0, *γ* = 0.9 min^-1^, *k*_fus_ = 1.0 min^-1^. The curve for *k*_fis_ = 0 (blue) corresponds to the de novo-fusion-decay. (B) Here, we vary the fusion rate *k*_fus_, while keeping *k_d_* = 0.7 min^-1^, *k*_fis_ = 0.8 min^-1^, *γ* = 1 min^-1^. The other values are the same. The curve for *k*_fus_ = 0 (blue) is included for comparison with the de novo synthesis-fission-decay model.

#### 4.1.2 Fission-fusion

Next, we explore the transient properties of the fission-fusion model. Biogenesis of mitochondria and vacuole is controlled through a balance of fission and fusion in yeast and mammalian cells [4, 21–24]. The steady-state organelle abundance for this model is given by a truncated Poisson distribution, which is characterized by the ratio of the fission and fusion rates. We seek to explore how organelle abundance evolves in time to attain steady state. To this end, we consider different values of fission and fusion rates and used a binomial distribution which was truncated at zero. The mean abundance increases monotonically to eventually reach the steady state value (see SI). In contrast, Fano factor behaves non-monotonically with time, often exhibiting a maximum, see Fig. 4 C, D. Interestingly, the transient Fano factor can exceed one when the rate of fission is higher than the fusion rate. This is a key result as, in the steady state, the Fano Factor for this process is bounded by one. This non-monotonic behavior diminishes as the average organelle number at *t* = 0 grows, even for the same parameter values (see SI). In order to estimate the effect of the spread of the initial distribution at *t* = 0, we choose binomial distributions with a fixed mean but different variances (see SI Fig. S2). There is a slight increase in the maximum Fano factor for a noisier initial distribution, but the behavior of the mean has very little dependence on the nature of the initial distribution and it monotonically reaches the steady-state value. Notably, while the steady-state mean and Fano factor depend on the ratio of fission and fusion rates, the transient behavior depends on the value of these individual parameters.

For the aforementioned models, consisting of two competing processes, the Fano factor clearly shows qualitatively distinct behavior as a function of time. In the ensuing section, we consider models with three processes.

#### 4.1.3 De novo synthesis-fission-decay

De novo synthesis-fission-decay model is often invoked to describe the biogenesis of peroxisome in yeast [4, 16]. While the steady state of this model is characterized by two effective parameters (ratio of de novo synthesis and fission rates, and fission and decay rates respectively), the transient behavior is more complex. Using the generating function technique [9], we compute the mean 〈*n*(*t*)〉 and Fano factor *F*(*t*) of organelle abundance as a function of time. The mean and Fano factor are given by

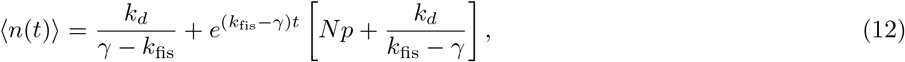

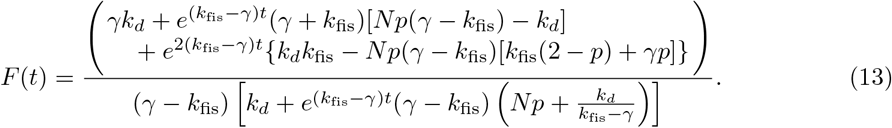

Using these expressions, we investigate the time evolution of the mean and Fano factor as the different parameters are tuned (see Fig. 3 A, B). As earlier, the mean abundance shows a monotonic behavior for all the parameter values. The trajectory of the mean with time depends exclusively on the mean of the initial distribution at *t* = 0 and does not show any dependence on the width of the distribution. The Fano factor, however, shows non-monotonic behavior and has a maximum for low values of the de novo synthesis rate. The steady-state Fano factor is independent of the de novo synthesis rate and the initial distribution, but the maximum Fano factor the system achieves is dependent on these parameters. This maximum is prominent for smaller values of the de novo synthesis rate (which results in a lower steady-state mean) and higher means of the initial distribution of organelle numbers (see SI Fig. S3). The behavior of the Fano factor has very little dependence on the width of the initial distribution. For higher values of the de novo synthesis rate or a lower mean of the initial distribution, the Fano factor behaves monotonically. As implied by the analytical solution for this system, the rate of convergence towards to steady state depends primarily on the difference between *k*_fis_ and *γ*.

#### 4.1.4 De novo synthesis-fusion-decay

The de novo synthesis-fission-fusion model is defined by three parameters, namely, de novo synthesis, decay and fusion rate constants). Like the previous model, we can expound the behavior of the mean and Fano factor as a function of time (see Fig. 3 C, D). The mean monotonically reaches the steady state value and its trajectory remains unchanged even if the variance of the initial distribution is altered. For certain values of the parameters, again a non-monotonic behavior is seen in the behavior of the Fano factor with time. Whether Fano factor exhibits a minimum or a maximum depends on the parameters used and the initial distribution. The maximum is seen when the degradation and fusion rates are low compared to the synthesis rate. Moreover, the maximum becomes more prominent with lower means in the initial distribution. The minimum is observed for higher values of the fusion rate and higher means in the initial distribution (see Fig. 3 D and SI Fig. S4). In most of these cases, the extrema are not very prominent compared to the other models. It must be noted that the introduction of fusion to the de novo synthesis-decay model qualitatively alters the behavior of the Fano factor even for a small fusion rate.

#### 4.1.5 De novo synthesis-fission-fusion-decay

Finally, we consider the general model consisting of all the four processes and explore how these four processes together impact organelle abundance distribution. As earlier, the mean abundance shows a monotonic behavior. The Fano factor for this model can exhibit a range of behavior, as one would expect owing to the increased complexity of the model. In fact, Fano factor may evolve monotonically or exhibit a minimum or a maximum as in the earlier models. The non-monotonic behavior seen in the Fano Factor for this system is short-lived compared to other models. The maximum may be seen for higher fission rates while the minimum is observed for fusion dominated processes. The maximum is more prominent for low organelle numbers in the initial distribution. Noise in the initial distribution does not affect the trajectory of the Fano factor by much, but higher noise may contribute to the height of the maximum if present. The trajectory of the mean has no influence of the noise (see SI Fig. S5).

## 5 Discussion

The abundance of organelles is highly regulated in cells. Unraveling the mechanism of organelle biogenesis has been a key area of investigation in cell biology. This area of research has received a boost in recent years owing to our ability to experimentally observe and count organelle copies at a single-cell level through live and fixed-cell imaging. The advent of these experimental techniques has provided a window of opportunity to undertake a theory-experiment dialogue in order to refine our understanding of the mechanisms of organelle biogenesis. In this manuscript, we focus on timelapse organelle abundance measurements, that remain largely untapped, to decipher how organelle abundance is regulated in response to internal and external cues. In particular, we theoretically investigate how the Fano factor of organelle distribution changes as a function of time for various mechanisms of organelle biogenesis. We find that different mechanisms exhibit qualitatively distinct behavior thereby allowing us to discriminate between these mechanisms.

Previous studies [4], that tried to falsify models of organelle biogenesis by comparing model predictions to data obtained from fixed cell imaging, were predicated on the assumption that organelle abundance distribution is in steady state. While this assumption has been useful in interpreting experimental data, it can lead to incorrect conclusions. For instance, the Fano factor for the fission-fusion model in the steady state never exceeds one, which was used as a unique feature of this model to discern it from other models [7]. In sharp contrast, the transient Fano factor for the fission-fusion model can actually be higher than one, as shown in Fig. 4C,D. Hence, it is imperative to be cautious while applying theoretical results to interpret experimentally measured organelle distributions, as obtained from fixed cell imaging. Moreover, the transient behavior of various models is richer than the steady-state behavior. For instance, the steady-state distributions of all the models we consider are functions of the ratio of the associated rate constants. As an example, the steady-state distribution of the fission-fusion model depends on the ratio of fission and fusion rate constants. The average organelle abundance can be increased by decreasing the fusion rate or increasing the fission rate. In steady-state, it is not possible to discriminate between these two possibilities. However, the time it takes to reach the steady state depends on which of the two rates is altered (see Fig. 4C,D). Hence, it is possible to discern between these two scenarios by observing the transient Fano factor.

The growing body of work on time-lapse measurements of organelle dynamics provides an alternative to fixed-cell imaging methods. For instance, recent experiments have reported real-time organelle dynamics in Drosophila [25], Hela cells [26, 27], Neurons [28]. These experiments contain information about organelle abundance; note that counting individual organelle copies from time-lapse videos is challenging [29]. By tapping into these datasets, and comparing them with the specific predictions various models make about the transient Fano factor, it should be possible to discriminate between them. We believe the results presented in this paper should pave the way for furthering the theory-experiment interplay in order to shed light on the process of organelle biogenesis.

## Acknowledgments

SC acknowledges the support provided by the DBT Ramalingaswami Fellowship.

## Supplementary Information

### 1 Transient solutions using the truncated ME

Eukaryotic cells tightly regulate organelle abundance in response to inter and intra-cellular cues. In this manuscript, we seek to obtain the transient properties of organelle abundance distribution. For the model of organelle biogenesis we consider (see Fig.1 in main text), we study the time evolution of organelle copy number distribution using a master equation approach. The probability *P_n_*(*t*) of having *n* organelle copies in a cell at time *t* is given by

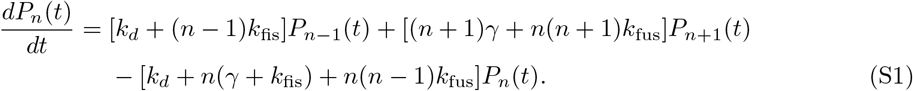

Owing to the nonlinear nature of Eq. S1, exact closed form expression for the time dependent probability distribution doesn’t exist. Here we employ an alternative route based on a recent paper that developed a method to obtain time dependent solutions for a general one-dimensional one-step master equation [8]. Eq. S1 is an infinite set of coupled differential equations, but in reality there will be an upper bound on the number of organelles. Taking this upper bound to be some large integer *τ*, we can truncate the master equation at *P*_*τ*–1_(*t*) and ignore all the higher terms. *P_n_*(*t*) can be written as a finite vector, *P*(*t*) = (*P*_0_(*t*), *P*_1_(*t*), *P*_2_(*t*),…, *P*_*τ*–1_(*t*))^*T*^. Hence, we can rewrite Eq. S1 as

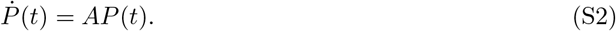

Here *A* is the transition matrix, given as

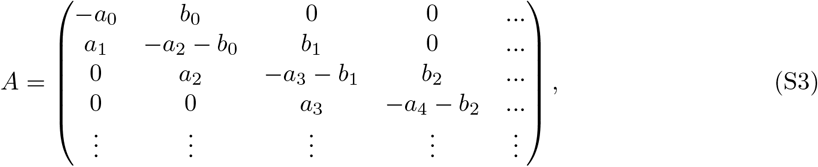

where *a_n_* = *k_d_* + (*n* – 1)*k*_fis_ and *b_n_* = (*n* + 1)*γ* + *n*(*n* + 1)*k*_fus_. The solution to Eq. S2 is

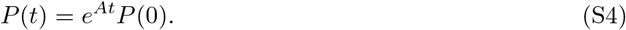

Solving this equation requires us to compute the eigenvalues and the eigenvectors of *A*, which is computationally expensive [30]. However, the matrix is tridiagonal and the sum of its the elements along each column is zero. This method, by reducing the problem to one of finding eigenvalues of a tridiagonal matrix, allows us to numerically compute the transient distributions. The transient probability distribution, evaluated in this manner, is given by

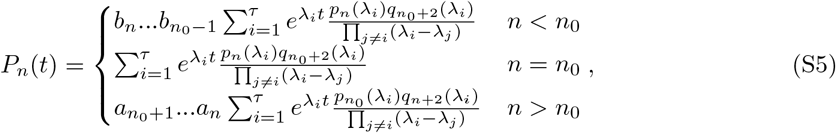

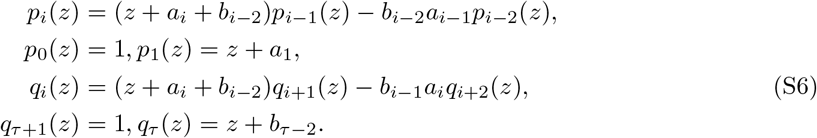

The solution given in Eq. S5 assumes that all the cells in the population have the same number of organelles, *n*_0_, at *t* = 0. This can be generalized to any initial distribution *P_n_*(0) = *Q_n_*, with the transient solutions now being given by

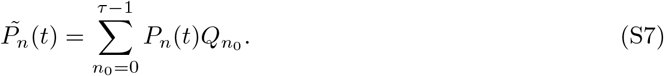

We assume two cases for the initial condition; a delta function distribution, i.e., all cells having the same organelle copy number, and a binomial distribution. These are expanded upon in further detail in the ensuing sections.

### 2 Exact time-dependent solutions for mean and variance of organelle copy number distribution

In the absence of fusion, Eq.S1 becomes a linear equation. Two of the models we consider belong to this category, namely i) De novo synthesis-decay and ii) De novo synthesis-fission-decay. We use the generating function technique to solve for the transient mean and variance for these aforementioned models [9].

#### S2.1 De novo synthesis-decay

Let us consider the case where only de novo synthesis and decay of organelles occur. The probability *P_n_*(*t*) of finding *n* organelle copies at time *t* is given by the master equation

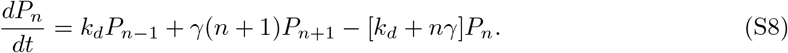

The generating function is defined as

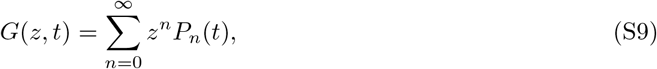

where 0 < *z* ≤ 1. We consider the derivatives of the *G*(*z, t*), as they will be important for the subsequent calculations.

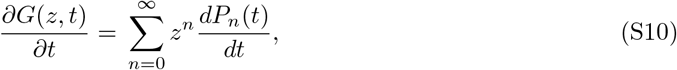

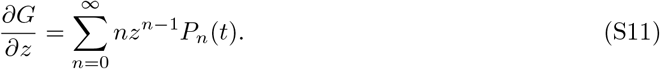

We multiply both sides of Eq. S8 by *z^n^* and sum over all possible values of *n* to obtain

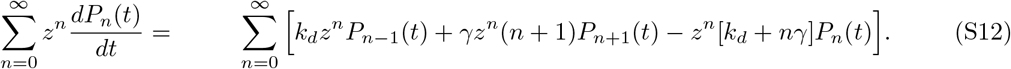

Using Eq. S10 and S11, we can rewrite Eq. S12 as

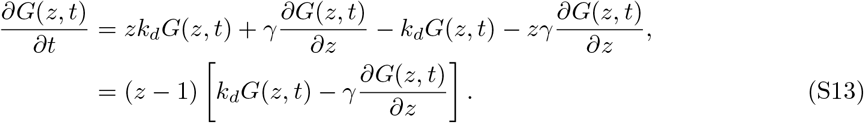

We get the following partial differential equation, which needs to be solved to solve for the generating function,

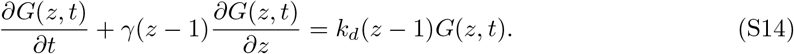

Eq. S14 can be solved analytically using the method of characteristics [10]. Using this method, we obtain the auxiliary equations of Eq. S14, given by

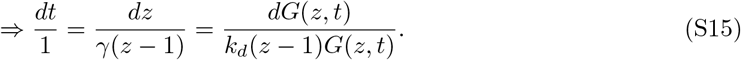

Integrating the first two terms leads to

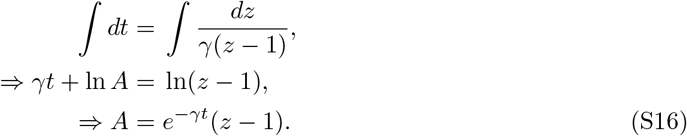

Here, *A* is the constant of integration. Similarly, integrating the second and third terms in Eq. S15 gives us

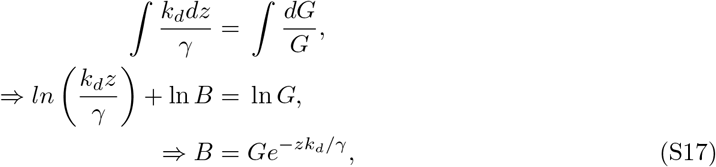

where *B* is the constant of integration. Writing *B* = *f*(*A*), where *f* is some function to be evaluated, we get

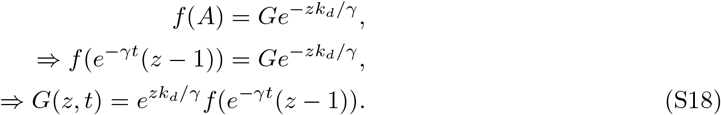

In order to evaluate the generating function, we need the initial conditions. Below, we consider two different initial conditions.

##### 2.1.1 Identical organelle copy number in every individual cell

Let us assume that at time *t* = 0 every cell in a population contains the same number of organelle copies, *N*_0_. In other words, the initial distribution has a delta function distribution *P_n_*(*t* = 0) = *δ*(*n* – *N*_0_), giving us the generating function at *t* = 0, *G*(*z*, 0) = *z*^*N*_0_^. Using this initial condition together with Eq. S18 gives us the functional form of *f*(*z*),

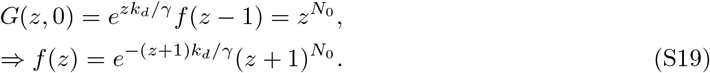

Using this form of *f*(*z*) in Eq. S18, leads to the following solution for the generating function,

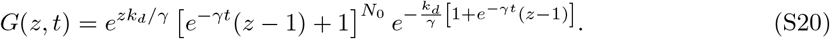

The generating function contains all the information about *P_n_*(*t*); the moments of *P_n_*(*t*) can be obtained from the generating function using the following relations

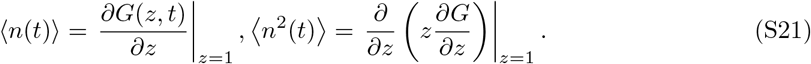

Using Eq. S21, we obtain the mean 〈*n*(*t*)〉 and the variance (defined as *σ*^2^ = 〈*n*^2^〉 – 〈*n*〉^2^),

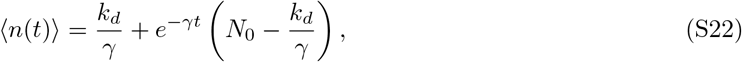

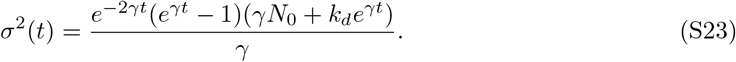

The Fano factor *F*(*t*), defined as the ratio of the variance to the mean is given by

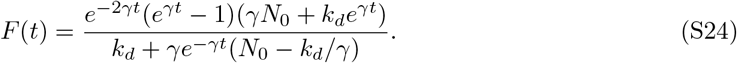

##### 2.1.2 Binomially distributed organelle copy numbers

We assume that the organelle copy numbers in individual cells follow a binomial distribution across a cell population, i.e. 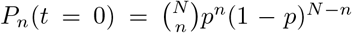. Here *N* is maximum number of organelle copies a cell can have and *p* is the corresponding probability for a cell to have a single organelle copy. From the definition of the generating function, at *t* = 0

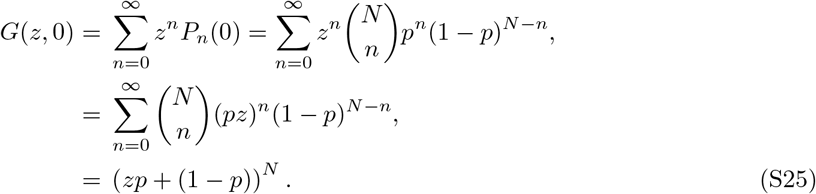

From Eq. S18 and Eq. S25, we get

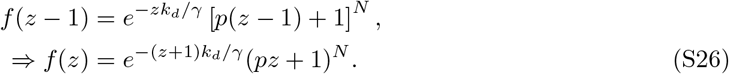

By substituting this form of *f*(*z*) in Eq. S18, we obtain an expression for the generating function

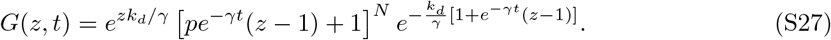

From Eq. S21 and S27, the mean and the variance are given by

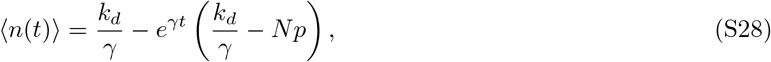

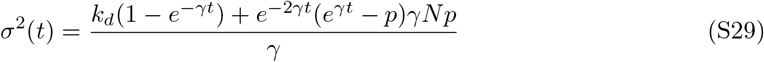

The Fano factor *F*(*t*) has the following form

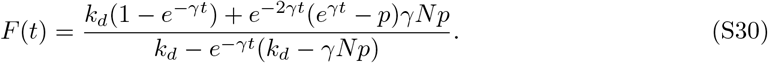

#### 2.2 De novo synthesis-fission-decay

Let us now consider a process in which the organelles undergo de novo synthesis, fission and decay. The probability for *n* organelles at time *t* obeys the following master equation,

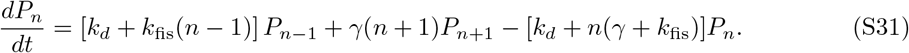

As in the previous section, we use the generating function, defined in Eq. S9. Multiplying both sides of Eq. S31 by *z^n^* and summing over all possible values of *n* gives us

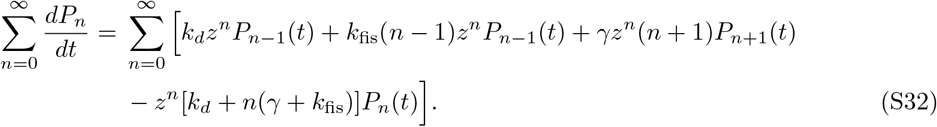

Using Eq. S10 and S11, we can rewrite Eq. S32 as

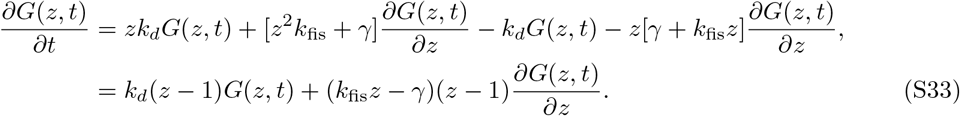

We get the following partial differential equation, which needs to be solved to obtain the generating function

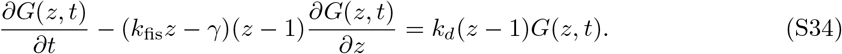

As before, we use the method of characteristics to solve Eq. S34. The auxillary equations of the partial differential equations are

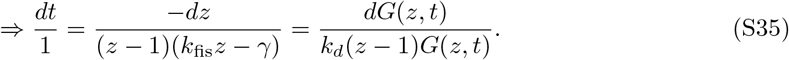

Integrating the first two terms gives

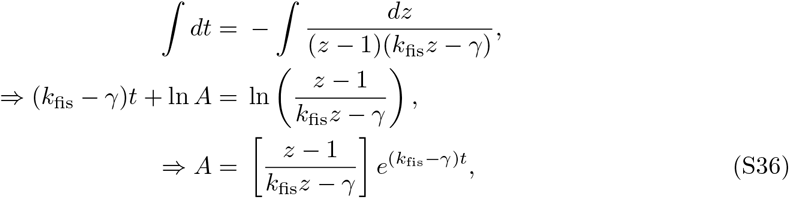

where A is the constant of integration. Similarly, integrating the second and third terms in Eq. S35

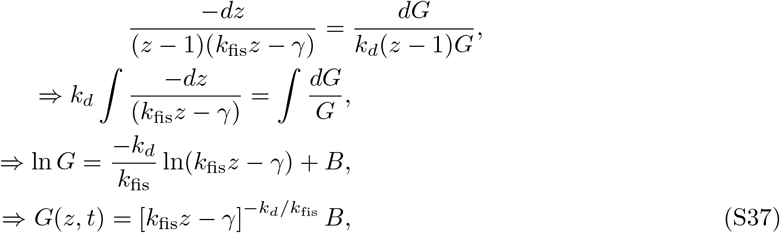

where *B* is the constant of integration. Writing *B* = *f*(*A*), where *f* is some function to be evaluated,

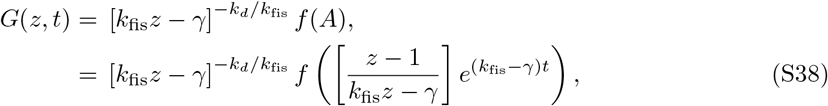

where we have used the expression for A from Eq. S36. Now, we will use the initial condition, and like previously, we consider two cases.

##### S2.2.1 Identical organelle copy number in every individual cell

We assume that initially all the cells have the same number of organelles *N*_0_. Then *P_n_*(*t* = 0) = *δ*(*n* – *N*_0_) and the generating function becomes *G*(*z*, 0) = *z*^*N*_0_^. Using this initial condition with Eq. S38, at *t* = 0, gives us

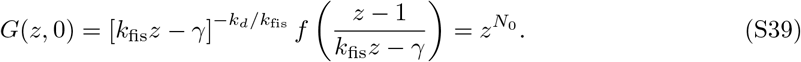

We make the following substitution into Eq. S39

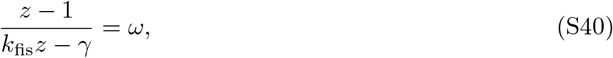

giving us

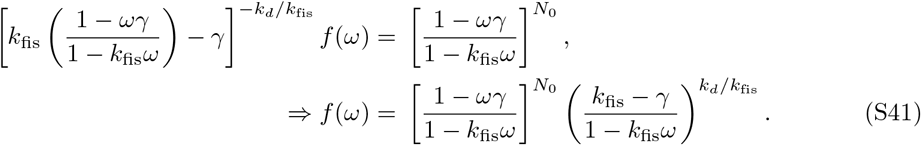

Now, we have the functional form of *f*(*z*). From Eq. S38 and Eq. S40, we get the generating function

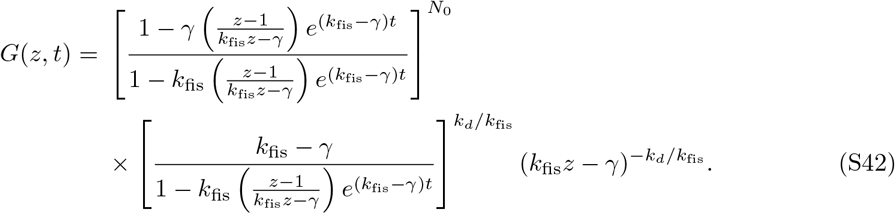

The mean and variance can be obtained using Eq. S21 and S42,

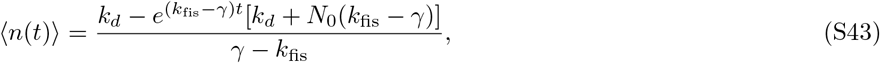

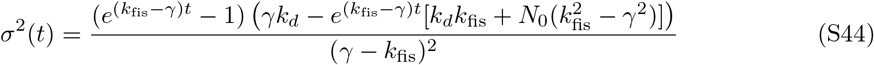

The Fano factor, *F*(*t*), has the form

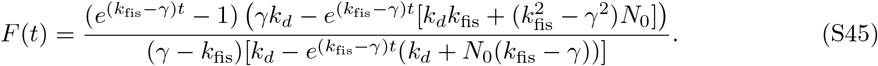

##### 2.2.2 Binomially distributed organelle copy numbers

We consider that the initial organelle abundance is binomially distributed i.e. 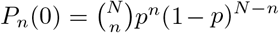. We get from the definition of the generating function

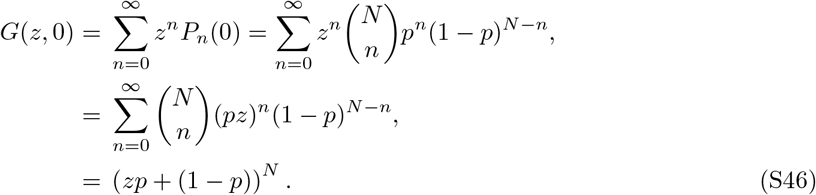

From Eq. S38 and S46, we get

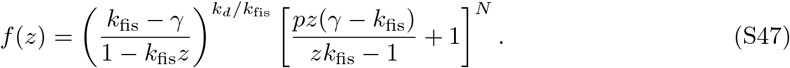

Substituting *f*(*z*) into Eq. S38, we obtain an expression for the generating function

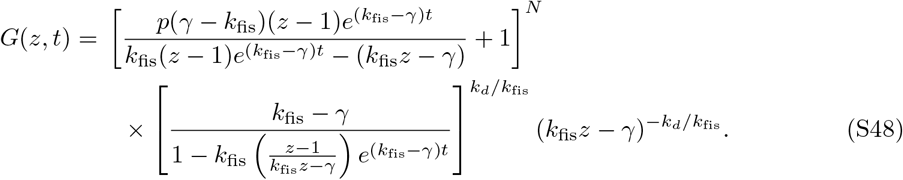

Using Eq. S21, the mean and variance are given as

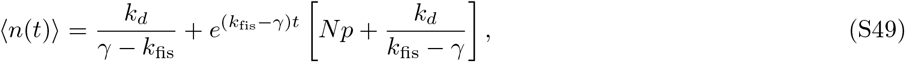

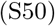

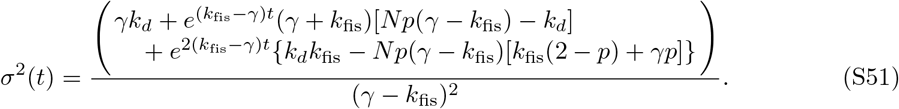

The Fano factor *F*(*t*) has the following form

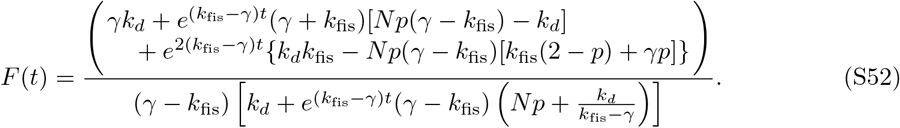

We plot Fano factor as a function of time for different models as the relevant model parameters are tuned.

### 3 Limitations of the model

Our model of organelle biogenesis makes a couple of simplifying assumptions. First, we assume that the rate constants associated with the various mechanisms of organelle biogenesis remain unaltered, and do not depend on organelle number or size. This may not always be the case [31, 32]). Second, although the de novo synthesis-fusion and de novo synthesis-fission-fusion models do have finite steady-states, the presence of the de novo synthesis process leads to ever increasing organelle biomass unless there are additional processes or regulatory mechanisms to account for it, thus these models have not been explored here.

### 4 Supplementary figures

**Figure S1:**
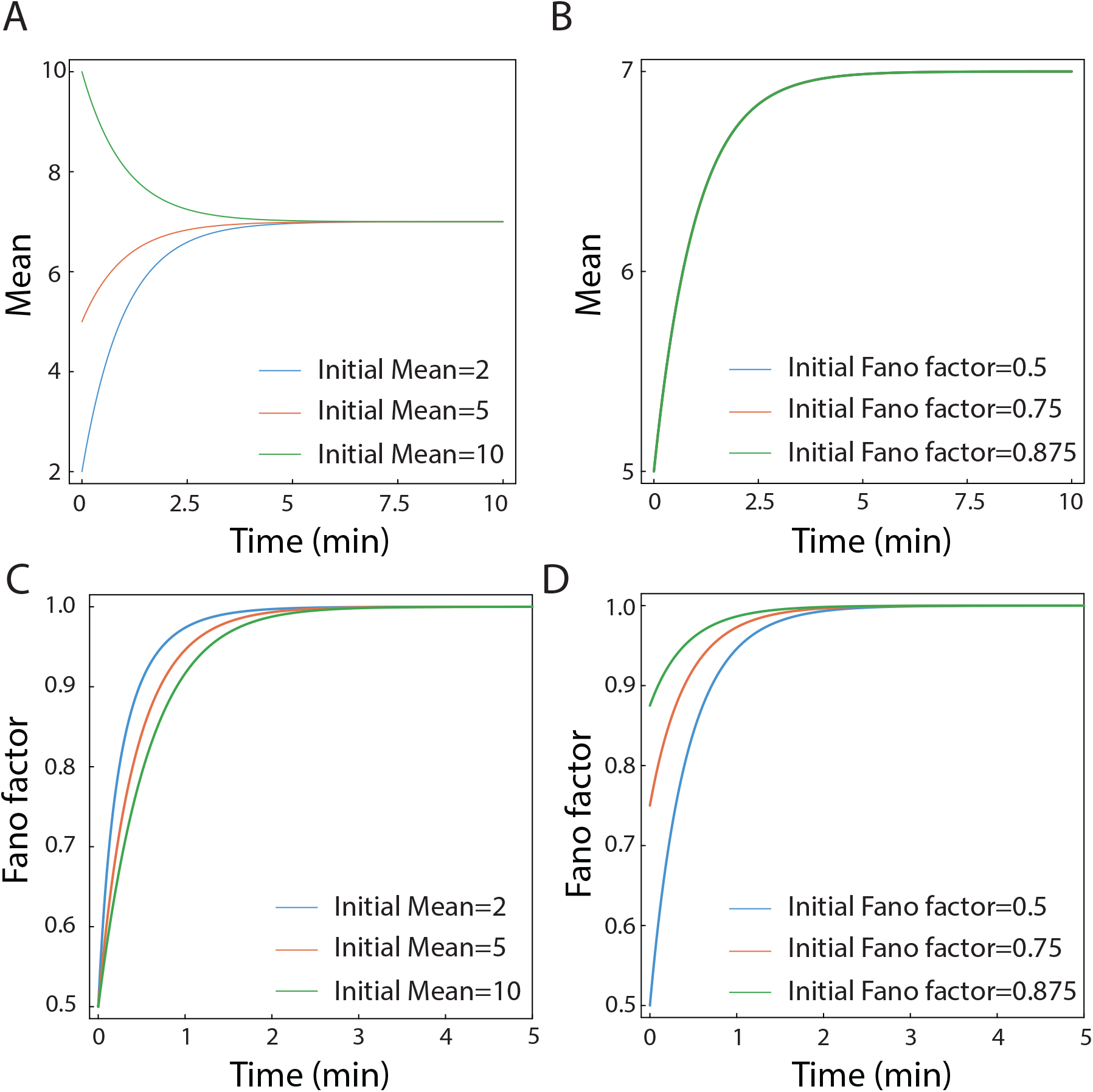
De-Novo Decay: (A) The evolution of the mean with time for different initial distributions, N = 30, 20, 10 p = 0.5. (B) The evolution of the mean starting from binomial distributions with mean = 5 and different variances, the trajectories overlap. (C) Evolution of the fano factor with time for different initial distributions as in (A). (D) The evolution of the fano factors with different initial distributions having the same initial mean, mean = 5, initial Fano = 0.5, 0.75, 0.875. *k_d_* = 7.0 min^-1^, *γ* =1.0 min^-1^ for all trajectories

**Figure S2:**
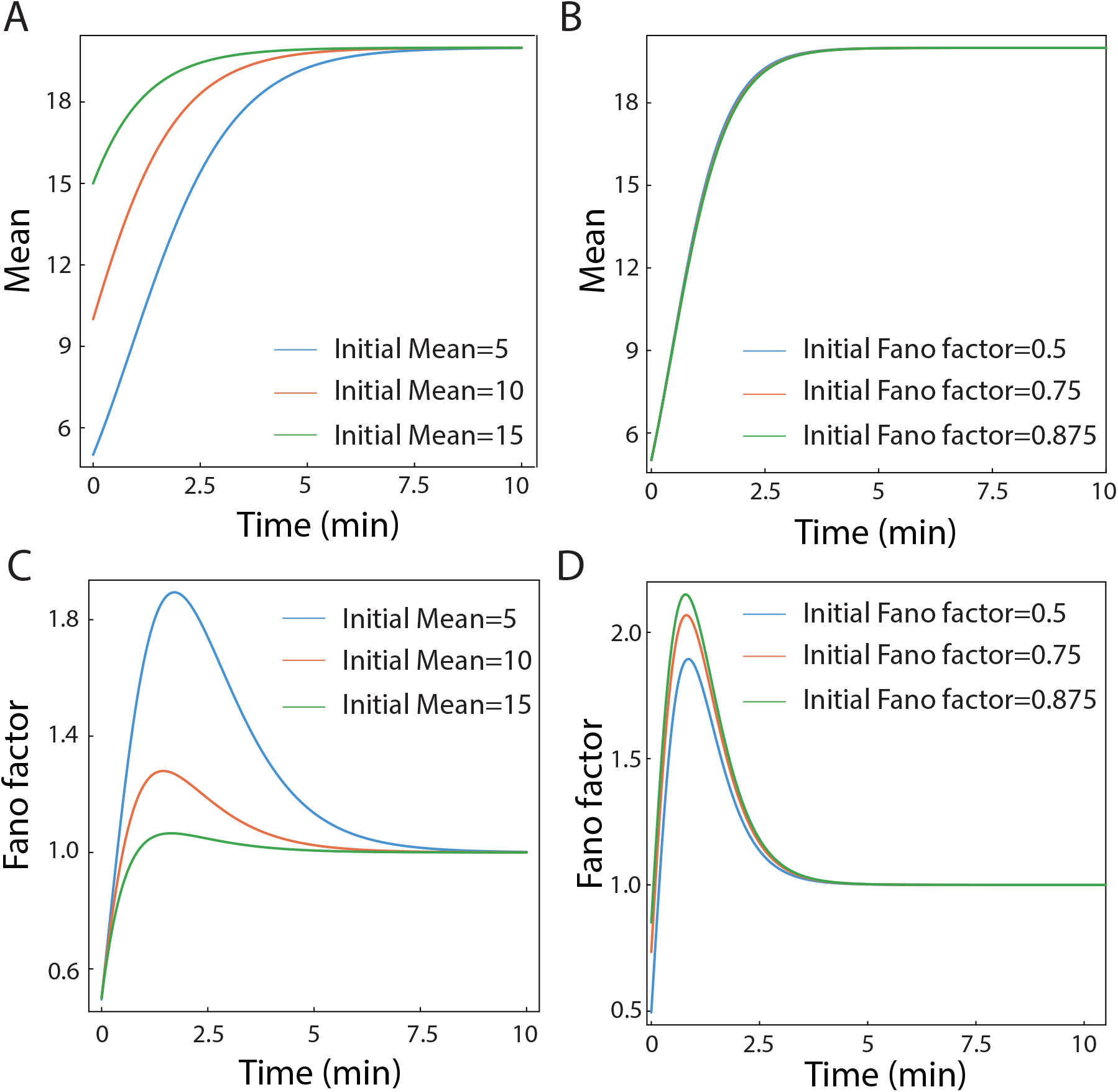
Fission - Fusion: (A) Evolution of mean when initial distributions of different means are chosen (blue = 5, red = 10, green = 15). (B) Evolution of the mean when different initial distributions with the same mean (5) but different Fano factors are taken (blue = 0.5, red = 0.75, green = 0.875). The trajectories overlap. (C) Evolution of the fano factor when initial distributions with different means are taken as in (A). (D) Evolution of the fano factor when different initial distributions are taken as in (B). *k*_fis_ = 1.0 min^-1^, *k*_fus_ = 0.05 min^-1^ for all trajectories.

**Figure S3:**
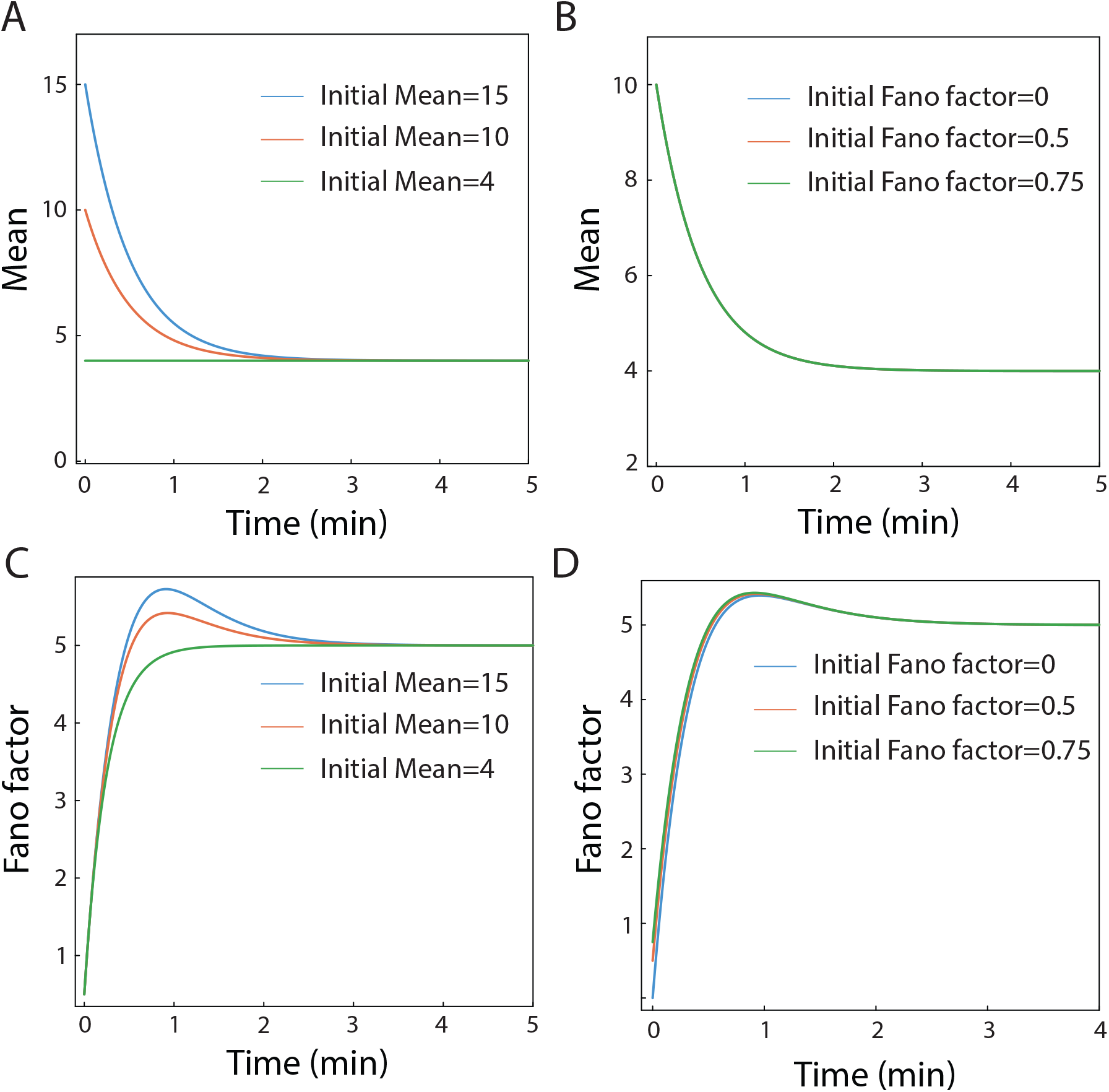
De novo - fission-decay (A) Evolution of mean when initial distributions of different means are chosen (blue = 4, red = 10, green = 15). (B) Evolution of the mean when different initial distributions with the same mean (10) but different Fano factors are taken (blue = 0.5, red = 0.75, green = 0.875). The trajectories overlap. (C) Evolution of the fano factor when initial distributions with different means are taken as in (A). (D) Evolution of the fano factor when different initial distributions are taken as in (B). *k*_fis_ = 8 min^-1^, *γ* = 10 min^-1^, *k_d_* = 8 min^-1^ for all trajectories.

**Figure S4:**
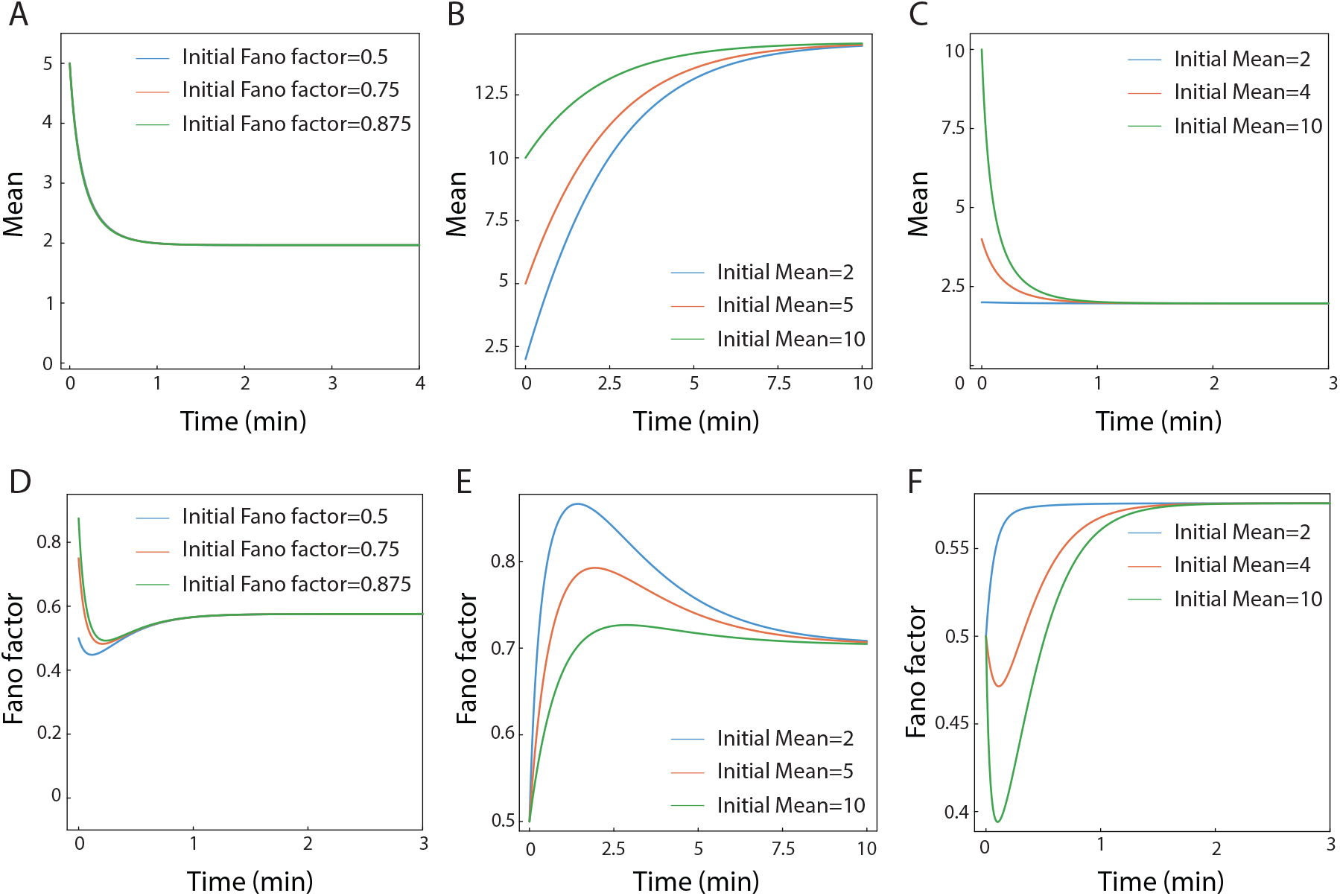
De novo-fusion-decay: (A) Evolution of the mean when initial distributions with the same mean (5) but different Fano noises are taken. *k*_fus_ = 1.0 min^-1^, *γ* = 1.0 min^-1^, *K_d_* = 5.0 min^-1^. (B),(C) Evolution of the mean when initial distributions with different means are taken. (B): *k*_fus_ = 0.01 min^-1^, *γ* = 0.2 min^-1^, *K_d_* = 5.0 min^-1^, (C): *k*_fus_ = 1.0 min^-1^, *γ* = 1.0 min^-1^, *K_d_* = 5.0 min^-1^. (D) Evolution of the Fano factor when distributions and parameters are taken as in (A). (E), (F) Evolution of the fano factor when the initial distributions and parameters are taken as in (B) and (C) respectively.

**Figure S5:**
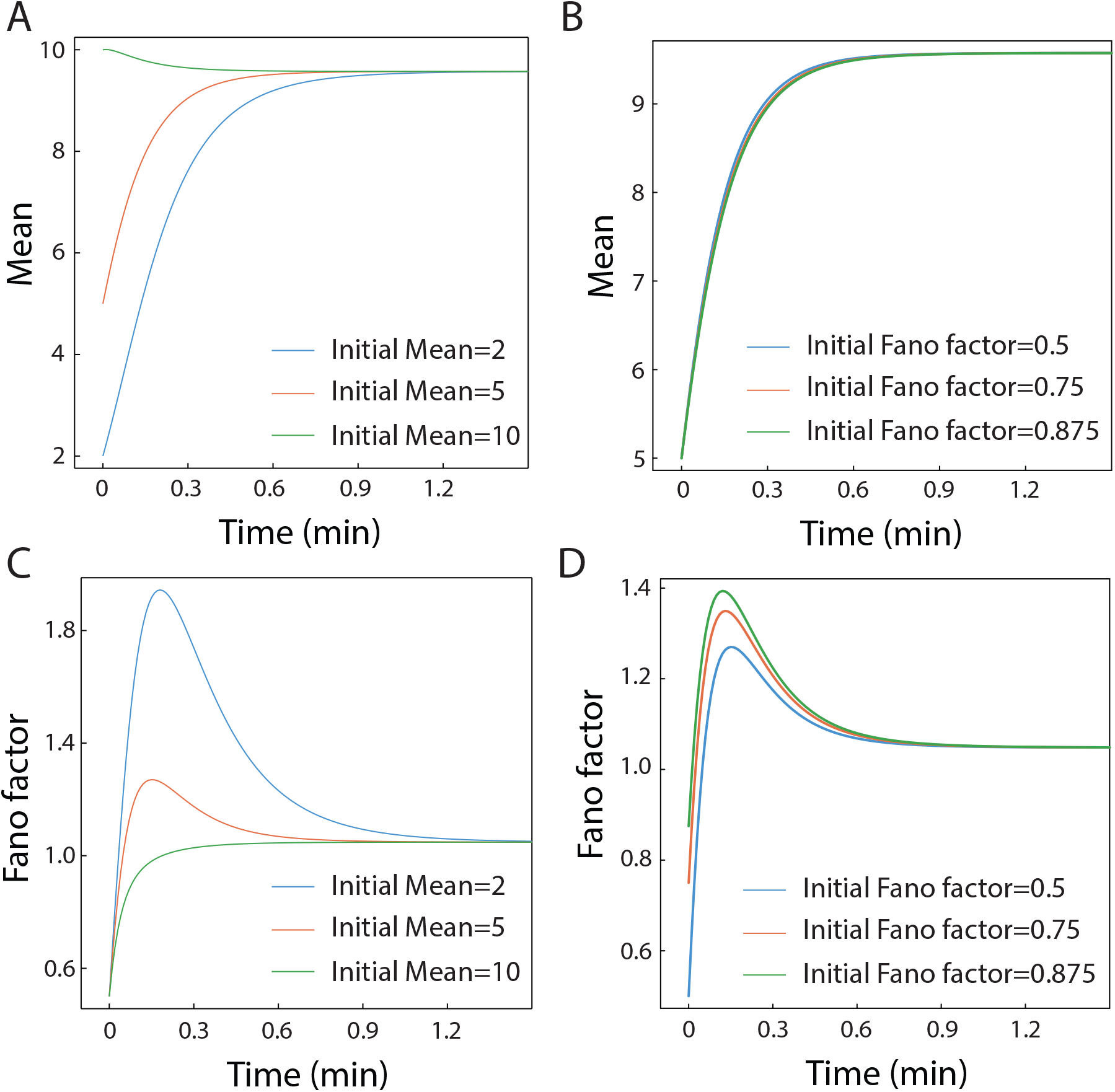
De novo - fission - fusion - decay: (A) Evolution of mean when initial distributions of different means are chosen (blue = 2, red = 5, green = 10). (B) Evolution of the mean when different initial distributions with the same mean (5) but different Fano factors are taken (blue = 0.5, red = 0.75, green = 0.875). The trajectories overlap. (C) Evolution of the fano factor when initial distributions with different means are taken as in (A). (D) Evolution of the fano factor when different initial distributions are taken as in (B). *k*_fus_ = 1.0 min^-1^, *k*_fis_ = 10 min^-1^, *γ* = 0.9 min^-1^, *K_d_* = 5.0 min^-1^ for all trajectories

## Notes

### Competing Interest Statement

The authors have declared no competing interest.

